# Lack of self-medication by fungus infected *Lasius platythorax* ants in a multitrophic experiment

**DOI:** 10.1101/2022.03.17.484742

**Authors:** Jason Rissanen, Heikki Helanterä, Torsten Will, Dalial Freitak

**Affiliations:** Institute of Biology at the University of Graz, Austria; Department of Ecology and Genetics, University of Oulu, Finland; Julius Kühn Institute (JKI) – Federal Research Centre for Cultivated Plants, Institute for Resistance Research and Stress Tolerance, Quedlinburg, Germany; Tvärminne Zoological Station, University of Helsinki, Hanko, Finland

**Keywords:** self-medication, ants, *Beauveria bassiana*, multi-trophic interactions, insect immunity, behaviour

## Abstract

Ants live in dense colonies with low genetic diversity between nestmates, creating favourable conditions for the spread of diseases. Due to the constant pressure by infectious disease, ants have evolved different mechanisms to cope with it. One way to combat pathogens is via consumption of biologically active compounds, which can inhibit the growth of the microbes or eliminate them. Although ants are known to respond to pathogen infections by altering their foraging to incorporate reactive oxygen species (ROS) into their diet to successfully combat disease under laboratory conditions, it is still unknown how ants would self-medicate in the wild. By designing a multitrophic environment, we try for the first time to identify potential sources of medicinal compounds for ants to use against fungal infections under natural conditions to better understand how the behaviour is expressed in nature. We investigated whether a bean plant suffering from an infestation of *Megoura viciae* aphids would be a source of ROS which the ant could use for self-medication, and whether the ant *Lasius platythorax* changes its foraging behaviour on bean plants if exposed to infectious spores of the entomopathogenic fungus *Beauveria bassiana*. We found no clear evidence that the ants significantly change their foraging behaviour on extrafloral nectaries in response to the pathogen or that the nectar produced by aphid stressed plants contains ROS. The aphids contained ROS in levels which ants could potentially use, however *Megoura viciae* is considered unpalatable to ants and we saw no evidence of predation on the aphids. Our results mean that the combination of species of the ant – plant – aphid interactions which we studied does result in a detectable self-medication response in ants. Therefore, we suggest that alternative species of aphids as well as other natural sources of ROS should be studied for self-medication.

## Introduction

Ants, like other social insects, live in dense colonies with a low genetic diversity between frequently interacting nestmates, making them susceptible for contracting and transmitting disease within their nests (Schmid-Hempel, 1998). Although there is evidence that pathogens are abundantly present both within and outside of their nest environments (Hughes et al., 2004; Valles et al., 2010; Reber & Chapuisat, 2012), colony collapse caused by infectious disease seems to be quite rare in nature (Evans, 1974). Apart from having physiological and behavioral strategies to help combat pathogens on an individual level, the interplay of individual defence strategies of ants can collectively mitigate the threat and effects of pathogens in their society as a form of social immunity (Cremer et al., 2007, 2018). However, social immunity does not make ant societies impregnable for disease. It is suggested, that social immunity is better equipped to limit the effects of generalist pathogens compared to specialist pathogens (Małagocka et al., 2019). Specialist pathogenic fungi like *Pandora* and *Ophiocordyceps* manipulate ant behavior to benefit their survival and transmission in ant colonies (de Bekker et al., 2014.; Małagocka et al., 2017; Araújo et al., 2018), but their occurrence in the natural environment of ants is considered to be rare compared to generalist pathogens, which are common in the surroundings of ant nests (Reber & Chapuisat, 2012) and thus predicted to have far greater effect on the colony fitness.

With the emergence of the social immunity theory, interest in pathogen caused changes in host behavior as beneficial for the host instead of for the survival and transmission of the pathogen has increased. Although the use of generalist entomopathogenic fungi in laboratory conditions has been questioned in terms of how relevant they are compared to natural conditions (Loreto & Hughes, 2016) there is a substantial amount of reported evidence that these fungi present a constant, even if not critical, threat to ants in nature (Hughes et al., 2004; Valles et al., 2010; Reber & Chapuisat, 2012). Due to the constant nature of the pathogen threat, it is likely that behaviour patterns of ants have adapted to deal with it through social immunity and individual behaviors. Behavioral changes in response to a pathogen infection which benefit ants has largely been studied with generalist entomopathogenic fungi such as *Metarhizium* and *Beauveria* (Cremer et al., 2018). In response to these pathogens, ants have been shown to regulate their rate of grooming (Tranter & Hughes, 2015) and allogrooming (Konrad et al., 2018), cut social ties and leave the nest if infected (Heinze & Walter, 2010; Bos et al., 2012), and alter their food intake to incorporate biologically active compounds as a form of self-medication (Bos et al., 2015; Rissanen et al., 2022).

Nutrition has recently been highlighted for special interest in the interaction between ants and pathogens (Csata & Dussutour, 2019). Immune responses to pathogen infections are costly in terms of energy and nutrients, and any deficiencies caused by disease can be offset through alteration in diet preferences as a form of compensatory diet choice (Ponton et al., 2011). Apart from merely replenishing diminished nutritional supplies, ants can use diet choices to directly combat pathogen infection through self-medication (Bos et al., 2015; Rissanen et al., 2022). Self-medication is the use of biologically active compounds in a way which would be harmful for uninfected individuals, and it can be therapeutic or prophylactic depending on when the compounds are used in relation to encountering the pathogen (de Roode et al., 2013; Abbott, 2014). Ants suffering from an infection by *Beauveria bassiana* successfully medicated themselves by incorporating food containing H_2_O_2_, a reactive oxygen species (ROS), under laboratory conditions (Bos et al., 2015), but the extent to which ants self-medicate and how they do it in nature in regards of obtaining and using biologically active compounds is still unclear.

Identifying self-medication behavior of ants under natural condition can be challenging (Sapolsky, 1994). As the pathogen threat for an ant nest is high it is likely that, if the ants are capable of self-medicating, it is incorporated in to their natural behavior (de Roode & Lefèvre, 2012). Therefore, changes in behavior caused by an active pathogen infection may be hard to detect. Compounds that insects have been shown to be able to use in a self-medication context have been studied as isolated compounds (Bos et al., 2015; Poyet et al., 2017) but their impact may be different when compared to their natural state inside e.g. specific plant organs. One of the criteria for self-medication suggested by de Roode et al., (2013) was, that the self-medication behavior must be relevant to the natural environment of the infected animals. Therefore, purely focusing on artificial foods in laboratory conditions is not sufficient to prove natural relevance, but instead there should be more focus on identifying natural pathways of medicinal compounds which ants could use.

Ants regularly enter symbiotic relationships with both plants (Rico-Gray & Oliveira, 2008) and aphids (Offenberg, 2001). Aphids offer ants honeydew, an excretion of excess sugars and liquids from the plant phloem on which they feed, in exchange for protection from predators (Offenberg, 2001). Regarding plants, some have extrafloral nectaries (EFNs) through which they reward ants with nectar for protection from herbivores (Rico-Gray & Oliveira, 2008). It is possible that ROS, produced systematically in response to e.g. aphid infestation – weak in the case of a compatible reaction and strong when an incompatible reaction between plant and a non-specialist species of aphids occur (Sun et al., 2020) - is also present in the nectar of infested plants, presenting ants with a natural source of ROS. Furthermore, EFN nectar contains a variety of other compounds, including antifungal compounds (Escalante-Pérez et al., 2012; Nepi, 2014) which ants could also potentially utilize. Aphids may act as a protein source in the case of need for increased protein as this can be a form of self-medication as well (Lee et al., 2006).

Here we study how an infection by the generalist entomopathogenic fungus *Beauveria bassiana* affects the foraging behavior of *Lasius platythorax* ants feeding on EFN bearing broad bean plants (*Vicia faba*) infested with aphids of the species *Megoura viciae. Lasius platythorax* is a common palearctic species of ants which generally builds its nest through excavating rotting wood (Seifert, 1991). *Megoura viciae* is a generalist pest on legume plants and does not engage in mutualistic behaviors with ants (Novgorodova, 2002) and is considered unpalatable or even potentially toxic to some insect predators (Tsaganou et al., 2004). The ants in our experiment have a quantitative feeding choice on foraging on the EFNs as well as a qualitative choice on consuming aphids. We are analyzing aphids for the presence of ROS as for apterous pea aphids (*Acyrthosiphon pisum*) a relevant concentration of hydrogen peroxide has previously described (Łukasik et al., 2012) to investigate whether aphids serve as a natural source of ROS that ants could use to self-medicate. We are analyzing ants for the presence of ROS to see, whether a change in foraging activity would correlate with changes in ROS concentrations in ants, possibly indicating a presence of ROS in the EFN nectar.

## Material and Methods

### Infection test

*Beauveria bassiana* was cultivated on petri dishes with Potato Glucose Agar and was incubated in the dark at room temperature. Spores were collected from plates with visibly sporulating fungus by pipetting 10mL of 1 × PBS directly on the plate and carefully rubbing it with a sterile glass rod. The spores were then centrifuged in 3000 rpm for 3 minutes and then suspended in 10mL of Milli-Q H_2_O. The spore concentration was quantified with a haemacytometer.

To assert the ability and timetable for which *B. bassiana* infects and kills the ants, we collected four wild nests of *L. platythorax* from tree stumps in a wood clearing in Lappohja, Hanko, Finland (59°54’46.9”N, 23°15’53.8”E). Each nest was split into four separate sub-colonies of 20 ants each, two of which were infected and the other two served as controls, for a total of 16 colonies (8+8). The colonies used in the infection treatment were infected by submerging the ants in a solution containing 1 × 10^7^ spores/mL of *B. bassiana* for five seconds. the control colonies were submerged in Milli-Q H_2_O for five seconds. Dead ants were counted and removed from the jars daily at 10 a.m. for a total of seven days. The colonies were all fed *ad libitum* with the Bhatkar & Whitcomb diet (Bhatkar & Whitcomb, 1970) and water.

### Experimental set-up

We collected 20 wild nests of *L. platythorax* from tree stumps in a wood clearing in Lappohja, Hanko, Finland (59°54’46.9”N, 23°15’53.8”E). Each nest was split into two sub-colonies of 500 workers each to be used in the two different treatments for a total of 40 colonies (20 colonies for infection treatment + 20 colonies for control treatment). The colonies used for the infection treatment were submerged in a solution containing 10^8^ spores/mL of *B. bassiana* for five seconds. The control colonies were submerged in Milli-Q H_2_O for five seconds. Each colony was then placed in individual plastic boxes (35 × 20cm and 20cm high) lined with a mixture of talcum powder and ethanol to keep the ants from escaping (Ning et al., 2019). Each box contained a 2cm layer of garden peat and a ceramic tile (10cm × 10cm) for nest-material.

Broad bean plants had been grown indoors in small plastic pots (6 × 6cm and 6cm high) in gardening soil, and the 40 plants in the best condition which had developed two pairs of EFNs were chosen for the experiment. 27 ± 3 *M. viciae* aphids were transferred from stock plants to the plants chosen for the experiment the day before the pots containing a plant and aphids were put into the nest-boxes with the ants. The variance in the number of aphids was to take the size differences of the individual aphids into account. The aphids were allowed to reproduce freely during the experiment.

### Foraging observation

The frequency of which the ants were visiting the EFNs in each colony was observed six times per day at 9 a.m., 11 a.m., 1 p.m., 3 p.m., 5 p.m., and 7 p.m. for a total of six days. During each observation period, the number of ants visiting a nectary was recorded. An ant was considered to be a forager if it was within a body length of an EFN and facing it. We also observed any behavior by the ants towards the aphids for signs that the ants would use the aphids as food, such as biting and carrying aphids into the nest.

### ROS analysis

Ants were sampled at the beginning of the experiment (day 0), the middle (day 3) and the morning after the end of the observational period (day 7). Three biological replicates (10 ants/sample) per each colony were taken randomly from the nest-box on each sampling event. Three biological replicates of aphids of mixed age were collected at the end of the experiment (15 aphids/sample). All the samples were placed in 250μl of 1 × PBS 3-amino-1,2,4-triazole solution (2 mg/mL) to prevent ROS from reacting, and stored in −80°C. The samples were homogenized with the TissueLyser II (Qiagen) for five minutes at 1800rpm. The samples were then centrifuged in 150 000rpm for 10 minutes in 4°C and the supernatant was collected for further analysis.

The protein content was analysed with the Bicinchonic Acid Protein Assay Kit (Sigma-Aldrich) according to the manufacturer’s protocol with minor adjustments, by using 12,5μl of sample in 100μl of working reagent. A dilution of 2,5μl of ant samples in 10μl of 1 × PBS was used for analyses. For the aphid samples, 5μl of the aphid sample in 7,5μl of 1 × PBS was used. The absorbance of each sample was read on 562nm wavelength in a microplate reader (SpectraMax iD3, Molecular Device).

The ROS content was analyzed with the Amplex Red Hydrogen Peroxide/Peroxidase Assay Kit (Invitrogen). The analysis was done according to the manufacturer’s protocol with minor adjustments, by using 25μl of sample in 25μl of reagent. A dilution of 12,5μl of ant samples in 12,5μl of reagent buffer was used. For the aphid samples 5μl of aphid sample in 20μl of reagent buffer was used. The fluorescence of the samples was read on 573 nm – 608 nm after excitation in 530 nm – 560 nm in a microplate reader (SpectraMax iD3, Molecular Device). The ROS/protein content was calculated to avoid differences caused by variation in ant and aphid sizes.

### Statistical analysis

All statistical analyses were performed with the R software (v 4.1.2).

The survival data of the infection test analyzed with a cox proportional hazard model by using the coxme function in the *coxme* package (Therneau, 2020). The day each individual ant died was used as the response variable with treatment used as a fixed factor and colony as a random effect to account for pseudoreplication.

The foraging data was analyzed with a poisson distribution general linear mixed model using the glmmTMB function from the *glmmTMB* package (Brooks et al., 2017). The number of foragers during an observational period was used as the response variable and the treatment as the explanatory variable. The original nest of the colonies was used as a random factor to account for any difference of the original nest in the analysis, and time was nested within colony as a random factor to account for pseudoreplication. Model diagnostics were analyzed using the *DHARMa* package for diagnostics of glmmTMB models (Hartig, 2018).

The ROS data was analyzed using the lmer function in the *lme4* package (Bates et al., 2015). ROS/protein was used as the response variable in the model with treatment as a fixed factor. The original nest was used as a random factor to limit any possible effects that they might have on the result.

Pairwise comparisons were done using the emmeans function from the *emmeans* package (Lenth, 2021) with a Tukey’s p-value adjustment when performing multiple comparisons.

Plants in two of the control colonies died during the experiment and therefore those colonies were omitted from all statistical analysis.

## Results

### Infection test

92.9% of all the ants who received the control treatment were still alive seven days after the treatment, whereas only 72.1% of the infected ants survived to that time. The effect of infection treatment on survival was significant (X^2^_1_ = 16.809, p < 0.001) (Figure 1). Most of the mortalities caused by the infection occurred during and after day 4.

**Figure 1.**
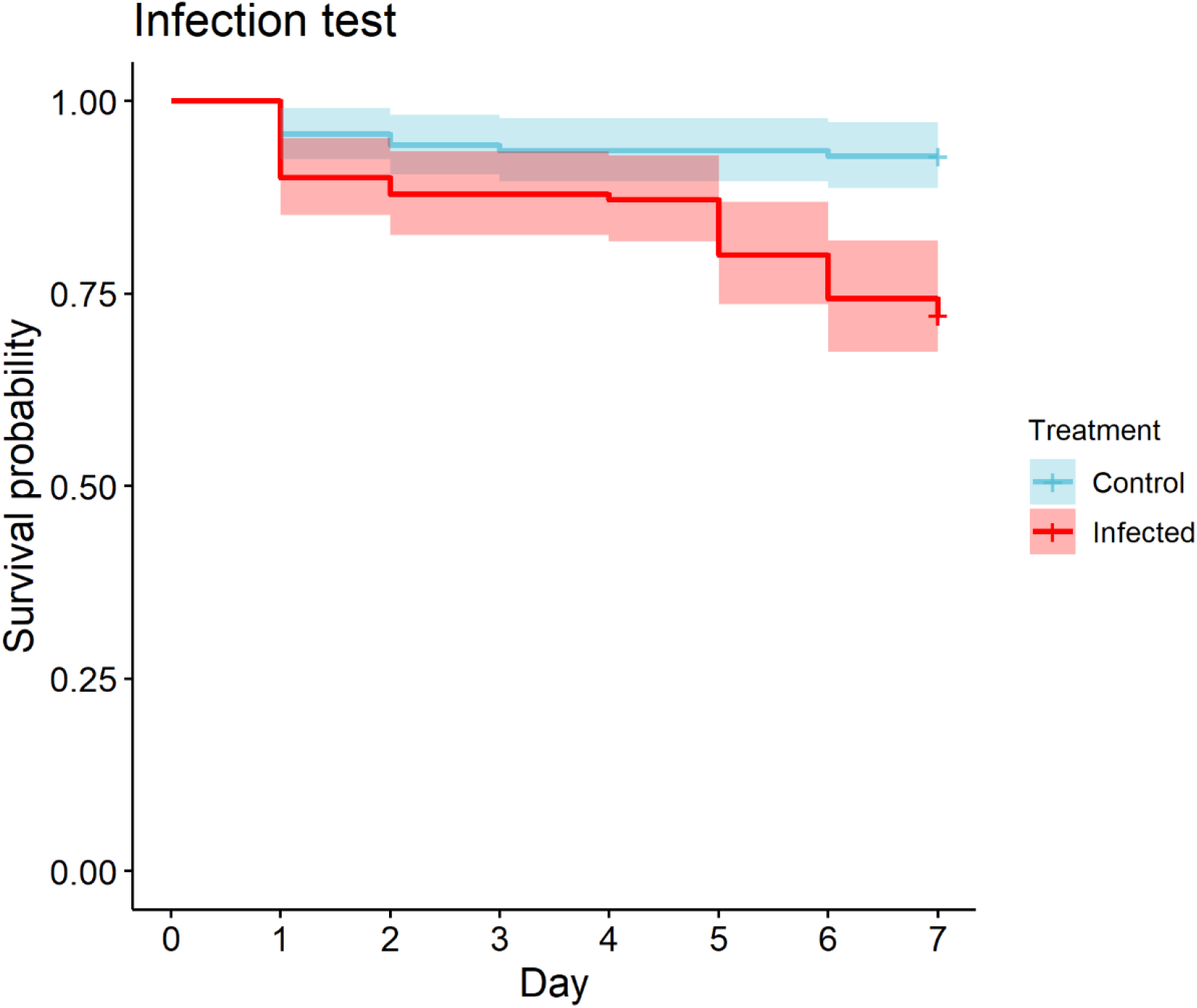
Survival graph of the infection test. The y-axis represents the probability of survival and the x-axis denotes time in days. *B. bassiana* was successful in infecting and killing workers of *L. platythorax* (X^2^_1_ = 21.15, p < 0.001). Most of the mortalities occurred after day 4. The red line indicates the survival of the infected colonies and the blue line represents the control colonies. The shaded area along the lines indicates the 95% confidence interval.

### Foraging results

As *B. bassiana* kills the majority of ants during and after the fourth day after infection, we decided to look at differences in foraging both before the infection kills the ants, the first four days after infection, as well as the foraging over the full six-day period.

Infection treatment did not affect the foraging frequency on nectaries in a significant way during the first four days (X^2^_1_ = 1.817, p = 0.178) (Figure 2a), nor did it affect foraging for the full six-day observation period (X^2^_1_ = 1.061, p = 0.303), Figure 2b).

**Figure 2.**
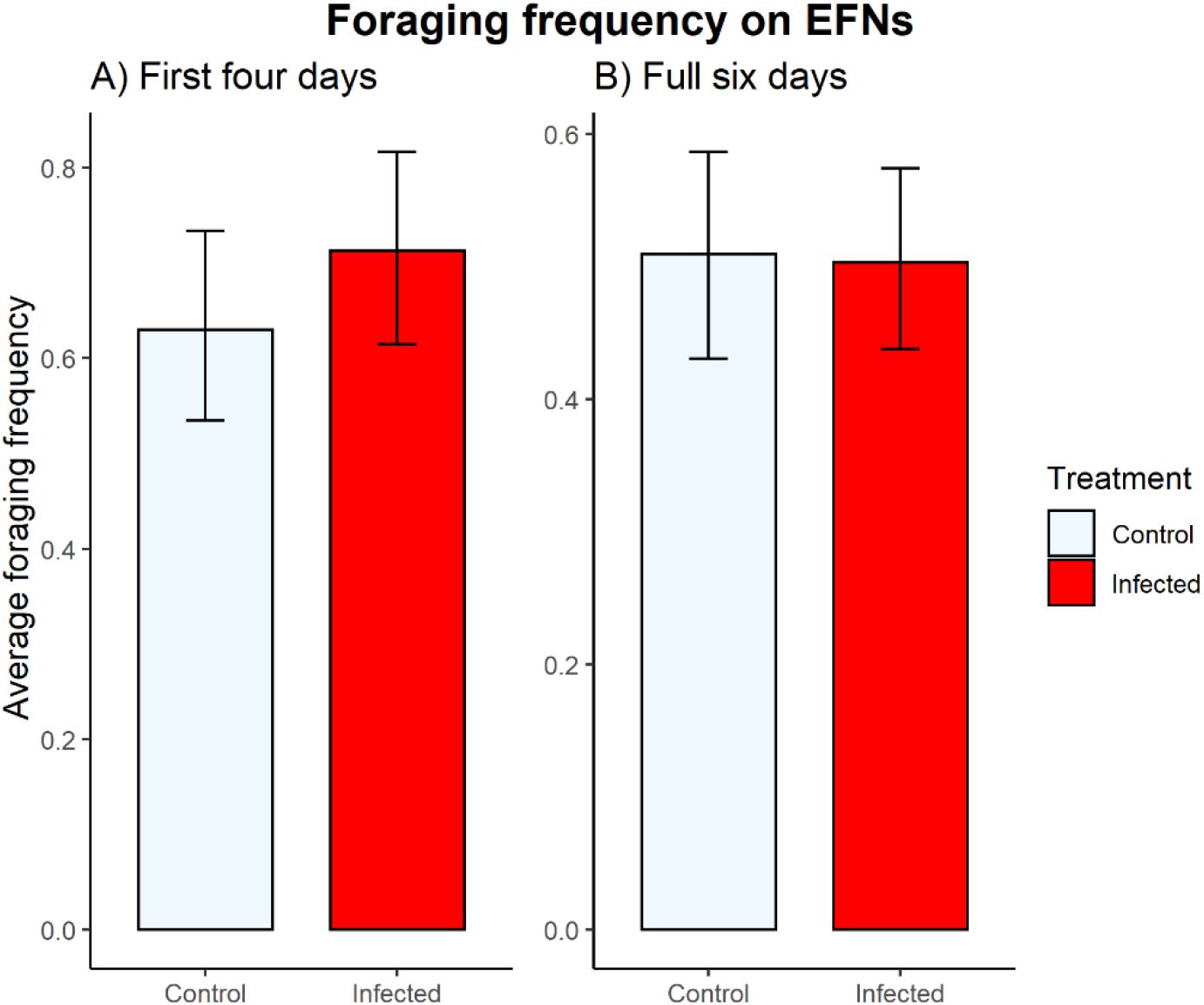
The foraging activity during the first three days (a) and the full 6-day observation period (b). The y-axis denotes the mean number of foragers found foraging at the EFNs at any time. No significant differences in foraging behavior was found between the infected (red) and control (black) colonies during either the first four days (t = −1.348, p = 0.178) or after the full six-day observation period (t = −1.030, p = 0.303). The error bars represent the 95% confidence interval.

### ROS analysis

The interaction of sampling day and infection treatment did not have a significant effect on the ROS content in ants (sampling day × infection treatment: X^2^_2_ = 3.786, p = 0.151). Neither the fixed effect of infection treatment (X^2^_1_ = 0.927, p = 0.336) nor the sampling day (X^2^_1_ = 5.190, p = 0.075) significantly affected the ROS content in ants. Within the infected colonies, ants had a higher concentration of ROS during the 7^th^ day compared to day 0 (t = −2.389, p = 0.046), but the difference was not significant between day 0 and 3 (t = −1.283, p = 0.406) or between day 3 and 7 (t=-1.106, p = 0.511). No differences in ROS concentration in the samples of healthy colonies were significant. Infected colonies contained a higher ROS concentration on day 7 compared to the control colonies (t = −2.124, p = 0.035). The aphids in our experiment contained ROS in a higher quantity compared to the ants on day 7 (t = −25.242, p < 0.001) (Figure 3).

**Figure 3.**
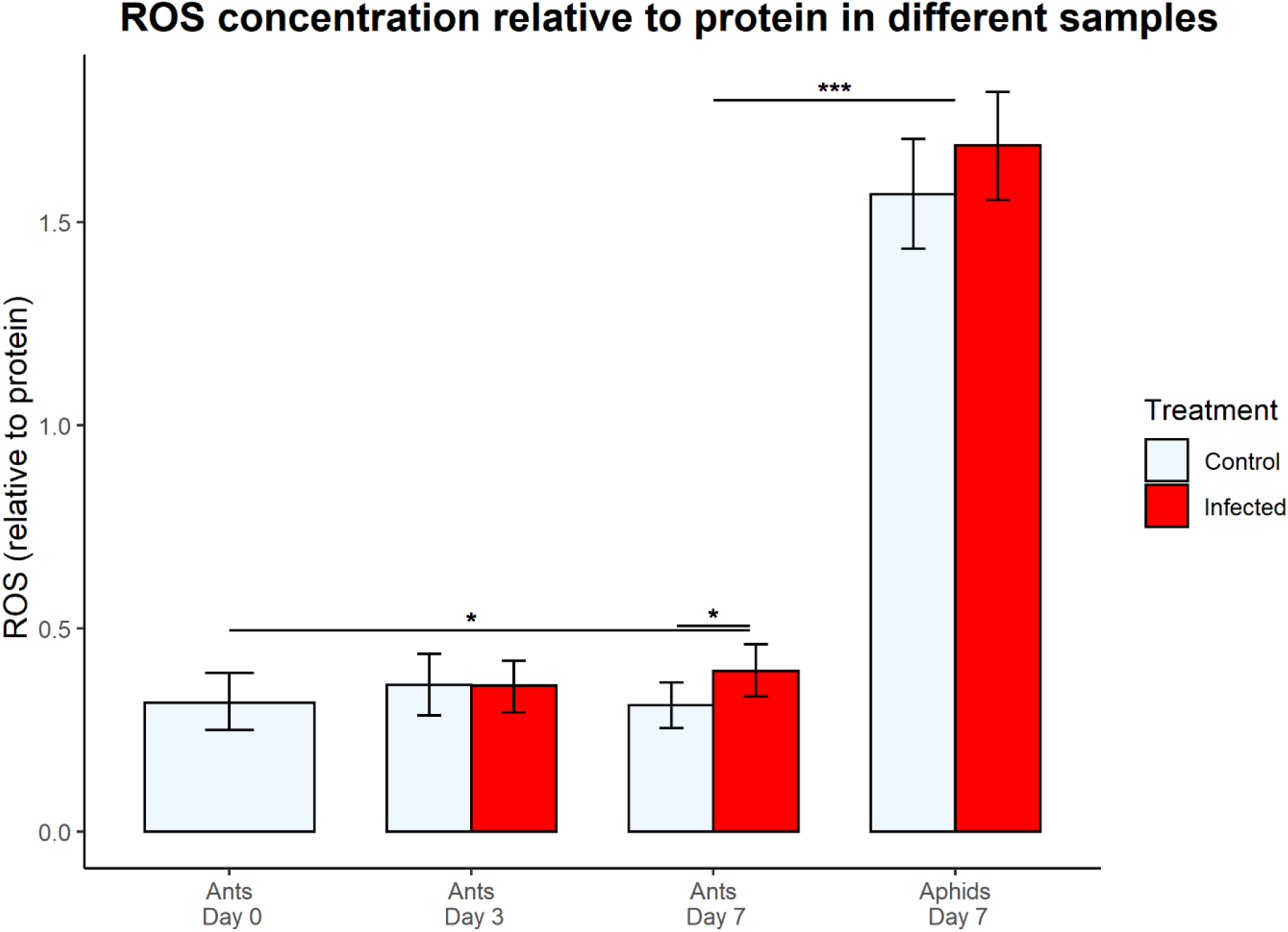
The differences in ROS content in mg/mL in each sample. The ants of the infected colonies had a significantly higher ROS content in the day 7 samples compared to the control colonies (t = −2.124, p = 0.035) and to the control at day 0 (t = −2.389, p = 0.046). The aphid samples contained a significantly higher ROS/protein content compared to the ants on day 7 (t = −25.242, p < 0.001). The error bars represent the 95% confidence interval.

## Discussion

Identifying natural pathways for ants to medicate themselves with is an area of research, which requires more focus. Whereas results of self-medication by ants so far have been obtained in laboratory conditions (Bos et al., 2015; Rissanen et al., 2022.), they are hardly representing natural environments which are significantly more complex. By using a multitrophic set-up, which reflects a natural environment for ants, we investigated whether infection by a fungal pathogen causes changes in the foraging behaviour of ants which could be deemed as self-medication.

In our experiment, we found no clear evidence that an infection of *L. platythorax* by *B. bassiana* elicits a significant change in the frequency of which *L. platythorax* ants visit the EFNs of an aphid infested plant. We did observe that ROS content in infected ants increase with time, whereas the ROS content in control ants was stable throughout. The increase ROS content in infected ants did not reflect the foraging activity of the ants, which in turn declined over time. The slight increase of ROS in infected colonies was most likely due to self-generation of ROS by the sick ants instead of ROS being present in the EFN nectar, which we were unable to directly examine in our experiment. This may be due to complex transport mechanisms in the plant because there is no close connection of glandular trichomes of the EFN and sieve tubes (Davis et al., 1988) potentially preventing ROS from entering the nectar even if the ROS response in the plant is systemic. Keeping the EFN nectar free of ROS during an aphid infestation could be beneficial for the plants in an evolutionary context, as the main purpose for a plant to invest in production of EFN nectar is to reward insect mutualists such as ants for protection against insect pests like aphids (Rico-Gray & Oliveira, 2008). If aphids would trigger a ROS response which is also expressed in the EFN nectar, it would reduce the quality of the nectar award for uninfected mutualists, therefore possibly reducing the protection behavior in exchange. EFN nectar can also contain a variety of other biologically active compounds (Nepi, 2014) as well as antifungal agents (Escalante-Pérez et al., 2012) which the ants could find attractive when infected.

Although no significant difference in foraging was observed as a response to the pathogen, the absence of self-medication cannot fully be claimed. It has been suggested, that when faced with a constant threat of a low-lethal disease, which *B. bassiana* is for ants in the wild (Reber & Chapuisat, 2012), evolution would favor that prophylactic medication would become a fixed behavior in ants (de Roode & Lefèvre, 2012). It is possible, that other compounds present in EFN nectar provides ants with medicinal benefits strong enough to combat low-level disease without a clear change in foraging behaviour. However, to prove that foraging on aphid stressed plants is a form of fixed prophylaxis and not compensatory feeding, the costs and benefits of consuming nectar from aphid stressed plants would have to be confirmed.

It is also important to consider the fact that we are only observing the foraging six times per day, but the feeding bouts of similar ants has been shown to last only up to two minutes (Portha et al., 2004), so it is possible, that even a small difference in the number of foragers when observed six times a day may accumulate to a bigger difference in nectar collection when considering the amount collected during the entire day. Therefore, an experiment with continuous observing of foraging activity would be needed to be able to properly prove or discredit any change in EFN nectar foraging. The ants in our experiment also only had access to one plant, whereas in nature a colony of ants would forage on several plants, accumulating the effect of any change in foraging due to a fungal infection on the total food intake of a colony.

The aphids in our experiment contained ROS in a higher quantity compared to the ants, presenting the ants with a potential source of both ROS and protein, both of which can be used for self-medicating (Lee et al., 2006; Bos et al., 2015). However, we found no supporting evidence for ants incorporating aphids into their diet. *Megoura viciae* is not an ant-affiliated species and there is no reported evidence that ants would use them as food, which could be due to their potentially toxic effects on some insect predators (Tsaganou et al., 2004). Our expectation was that ants would overlook the potential toxic effects of consuming the aphids due to their protein and ROS contents, yet this was not observed. However, our observational periods were limited, and it could be that the ants were in fact supplementing their diet with aphids, but we were not able to see it without constant monitoring. There is also the possibility that the ants would not shift their interest towards the aphids if they found the compounds they needed in the nectar.

The results of this study show, that an infection by a generalist entomopathogenic fungus does not induce a clear change in foraging behavior in ants on a plant infested with *M. viciae* aphids. Discovering the extent of self-medication behaviour in ants and especially how they do it in the wild remain important questions to answer. There is a possibility, that using palatable aphids could provide the ants with a more usable source of ROS, if they contain it in the same level as *M. viciae* in our experiment. It is also possible, that the ant – plant – aphid interaction does not present the ants with a viable source of medicine, and instead experiments using alternative sources of biologically active compounds available for ants should be pursued.

